# Can non-invasive brain stimulation modulate peak alpha frequency in the human brain? A systematic review and meta-analysis

**DOI:** 10.1101/2023.11.13.566909

**Authors:** S.K. Millard, D.B. Speis, P. Skippen, A.K.I. Chiang, W.J. Chang, A.J. Lin, A.J. Furman, A. Mazaheri, D.A. Seminowicz, S.M. Schabrun

## Abstract

Peak alpha frequency, the dominant oscillatory frequency within the alpha range (8–12 Hz), is associated with cognitive function and several neurological conditions, including chronic pain. Manipulating PAF could offer valuable insight into the relationship between PAF and various functions and conditions and provide new treatment avenues. This systematic review aimed to comprehensively synthesise effects of non-invasive brain stimulation (NIBS) on PAF speed. Relevant studies assessing PAF pre- and post-NIBS in healthy adults were identified through systematic searches of electronic databases (Embase, PubMed, PsychINFO, Scopus, The Cochrane Library) and trial registers. The Cochrane risk-of-bias tool was employed for assessing study quality. Quantitative analysis was conducted through pairwise meta-analysis when possible; otherwise, qualitative synthesis was performed. The review protocol was registered with PROSPERO (CRD42020190512) and the Open Science Framework (https://osf.io/2yaxz/). Eleven NIBS studies were included, all with a low risk-of-bias, comprising seven transcranial alternating current stimulation (tACS), three repetitive transcranial magnetic stimulation (rTMS), and one transcranial direct current stimulation (tDCS) study. Meta-analysis of active tACS conditions (eight conditions from five studies) revealed no significant effects on PAF (mean difference [MD] = -0.12, 95% CI = -0.32 to 0.08, p = 0.24). Qualitative synthesis provided no evidence that tDCS altered PAF and moderate evidence for transient increases in PAF with 10 Hz rTMS. There is limited evidence that NIBS can modulate PAF, and existing evidence does not demonstrate robust effects. Further studies employing standardised stimulation protocols are necessary to determine the potential of NIBS to modulate PAF.

## 1 Introduction

Alpha is the dominant oscillatory frequency (8–12 Hz) recorded in the human brain using electroencephalography (EEG) or magnetoencephalography (MEG) (Van Diepen et al., 2019). The frequency exhibiting the highest power within the alpha range, termed the peak alpha frequency (PAF) or individual alpha frequency (IAF), is relatively stable within individuals (Kondacs and Szabó, 1999) and possesses a trait-like quality, with heritability accounting for a significant portion (71–83%) of the variance in PAF across individuals (Posthuma et al., 2001). Faster PAF (i.e. higher frequency) is associated with improved cognitive performance in working and semantic memory tasks (Klimesch, 1999). PAF is also correlated with individual processing capacity, both in trait (i.e. inter-individual) and state (i.e. intra-individual) contexts (Minami and Amano, 2017). PAF follows a developmental trajectory, increasing throughout childhood, stabilising in late-adolescence/adulthood (∼10 Hz), and decreasing in old age, effectively paralleling age-related changes in brain volume and cognitive performance (Bigler et al., 1995; Breslau et al., 1989).

Individuals with depression (Tement et al., 2016), post-traumatic stress disorder (Wahbeh and Oken, 2013), autism (Dickinson et al., 2018), and chronic pain (Fauchon et al., 2022; Kim et al., 2019; Sarnthein et al., 2006; Vries et al., 2013) exhibit slower PAF than the general population. Interventions that can modulate PAF may be useful to help illuminate the role of oscillations for brain function in health and disease and potentially underpin new therapeutic approaches for a variety of conditions (Herrmann et al., 2013; Sejnowski and Paulsen, 2006). A number of interventions are thought to be capable of modulating brain oscillations, including non-invasive brain stimulation (NIBS).

NIBS techniques are a collection of safe technologies used to explore and modify brain activity without requiring invasive procedures like surgery (Herrmann et al., 2013). The primary NIBS techniques are transcranial magnetic stimulation (TMS), which applies a magnetic field to the brain via a coil positioned on the scalp, and transcranial electrical stimulation (tES), which applies weak electrical currents to the brain via scalp electrodes. The generated magnetic or electrical fields can penetrate the skull and interact with electrical fields produced by neuronal populations in the brain (Herrmann et al., 2013; Ridding and Rothwell, 2007). Through control of stimulation parameters (e.g., amplitude, frequency, duration) NIBS techniques can modify neural excitability, plasticity, and oscillatory activity in targeted brain regions (Bergmann and Hartwigsen, 2021; Braga et al., 2021; Herrmann et al., 2013; Mansouri et al., 2018; Polanía et al., 2018; Vogeti et al., 2022). Specifically, as oscillations reflect fluctuations in the electrical activity of neural populations over time (Biasiucci et al., 2019; Cohen, 2017a; Lopes da Silva, 2023, 2013; Nunez and Srinivasan, 2006) and NIBS directly alters electrical activity of neural populations (Bergmann and Hartwigsen, 2021; Braga et al., 2021; Herrmann et al., 2016, 2013; Mansouri et al., 2018; Polanía et al., 2018; Vogeti et al., 2022), NIBS techniques are prime candidates for PAF modulation. However, the existing literature lacks a systematic review of NIBS effects on PAF speed.

As PAF is closely associated with various cognitive functions and diseases, establishing whether NIBS interventions can modulate PAF could provide new avenues to influence and investigate brain function. The aim of this systematic review was to comprehensively synthesise the evidence for effects of NIBS interventions on PAF in healthy participants. Specifically, we aimed to determine: 1) the types of NIBS interventions used to modulate resting state PAF in healthy adults; 2) the magnitude, direction, and duration of any effects of NIBS on PAF; and 3) the sample sizes, methodological approaches, and environmental characteristics of studies that successfully modulated PAF.

## 2 Materials and methods

### 2.1 Protocol and registration

This systematic review was reported according to the Preferred Reporting Items for Systematic Reviews and Meta-Analyses (PRISMA) guidelines (Liberati et al., 2009; Moher et al., 2009). The protocol of this review was registered at the International Prospective Register of Systematic Reviews (PROSPERO; registration number: CRD42020190512) and has been made available on the Open Science Framework (OSF; https://osf.io/2yaxz/). Note that the initial protocol sought to explore the effect of a range of different interventions (e.g., NIBS, exercise, drugs) on PAF. Due to the heterogenous mechanisms of action and wide variety of interventions used to modulate PAF, this study only reports the effects of NIBS interventions. This protocol deviation was recorded on OSF (https://osf.io/2yaxz/).

### 2.2 Search strategy

Searches were conducted to find completed studies since 2000 in the following databases: EMBASE, PsychINFO, PubMed, Scopus, and the Cochrane Library. Search terms consisted of combinations of key terms referring to PAF, EEG, and neuromodulation interventions, using boolean operators and truncations to ensure sensitivity and specificity. The project team created exact search strategies with guidance from an expert librarian and adapted them for each database (Supplementary Material 1).

Trial registers and repositories of results, including the U.S. National Library of Medicine (https://clinicaltrials.gov/), the System for Information on Grey Literature in Europe (https://opengrey.eu/), the New York Academy of Medicine Grey Literature Report (http://www.greylit.org), and the Open Science Framework Preprint archive search (https://osf.io/preprints/discover) were searched to identify completed unpublished studies.

Database searches were carried out by the first author (SKM). The initial search, including all neuromodulation techniques, was conducted in February 2021, and a final repeat search of only NIBS interventions was conducted in April 2023 prior to publication of review outcomes. References of relevant articles and reviews were also searched manually for additional articles.

### 2.3 Inclusion criteria

1. Studies written in English;
2. Studies published after or in the year 2000;
3. Participants: healthy adults aged between 18 and 65 years, no restrictions on sex, gender, or race/ethnicity. Studies involving clinical populations that also assessed a healthy control group were included. Only information from the healthy control group was extracted;
4. Intervention: any intervention using NIBS techniques, including:
  i. magnetic stimulation techniques (e.g., repetitive transcranial magnetic stimulation [rTMS], theta burst stimulation [TBS]),
  ii. electrical stimulation techniques (e.g., transcranial alternating current stimulation [tACS], transcranial direct current stimulation [tDCS], cranial electrotherapy stimulation [CES], reduced impedance non-invasive cortical electrostimulation [RINCE], or transcranial random noise stimulation [tRNS]);
5. Comparison: studies with or without control groups were included;
6. Outcome: change in resting state PAF (Hz), with PAF measured using resting state EEG or MEG before and after an intervention. Resting state may include a relaxed supine, seated, or standing position with eyes either opened or closed;
7. Study design: original experimental or quasi-experimental research studies, using single group, parallel, or cross-over study designs with both randomised and nonrandomised allocation.

### 2.4 Exclusion criteria

1. Studies investigating patient populations (i.e. defined as registered to receive or receiving medical treatment) without a healthy control group;
2. Studies investigating populations other than humans (i.e., animal models, simulations, computer models);
3. Studies measuring EEG or MEG in a state other than conscious awake states (e.g., sleep, coma, vegetative state);
4. Studies only measuring PAF at one time point or during stimulation;
5. Reviews, commentaries, editorials, study protocols, conference abstracts or proceedings, book chapters, letters to the editor, or case studies.

### 2.5 Study selection

Search results were exported to Mendeley version 1.19 (London, UK), where duplicate articles were identified and removed. Two independent reviewers assessed titles and abstracts to identify potentially relevant studies. Any cases of doubt were automatically selected for full-text eligibility evaluation. The full-texts of these studies were retrieved and an automatic full-text scan phase was used to identify articles that reported measurement of PAF in the full text. A conceptual flow chart, key words, and the Python code used for the FT-scan are freely available (https://github.com/sammymillard/ft-scan). Full-texts were assessed by two independent reviewers against eligibility criteria. A third reviewer was consulted to resolve any disagreements. Excluded studies and the reason for exclusion were recorded.

### 2.6 Data extraction

A customised data extraction form was used by two independent reviewers to extract data from each relevant study. Any inconsistencies were resolved by a third reviewer. The following data were extracted:

- Study characteristics: study design, randomisation procedures, and number of conditions/groups in each experiment.
- Participant characteristics: sample size, sex, age, and any other demographic information provided.
- Interventions: exact NIBS intervention implemented, route of delivery, dose, duration, frequency, timing of intervention, as well as comparison conditions and co-interventions used.
- EEG recording: electrode numbers and location, eyes opened/closed, sampling rate, filter properties, room characteristics, participant position, duration of recording, timing of recording in respect to intervention.
- PAF calculation: length of epochs, frequency conversion method, bins, and range used, as well as PAF calculation method, and regions of interest.
- Outcomes: pre- and post-intervention PAF means and standard deviations (SDs), mean differences (MDs) and SDs, standardised mean difference (i.e. effect sizes), their variance and standard error (SE) of variance, as well as F-values, t-values, and p-values when means and SDs were not available.

### 2.7 Missing data

Corresponding authors were contacted up to three times via email to request missing data. In cases where no reply was received within six weeks of the third contact attempt, data were deemed inaccessible. Data were extracted from figures where possible.

### 2.8 Risk of bias and EEG methodological assessment

Risk of bias was assessed using version 6 of the Cochrane Collaboration’s tool for assessing risk of bias (Higgins et al., 2011; Sterne et al., 2019). Elements of methodology and reporting were assessed using the best practice recommendations for MEG and EEG (i.e. MEEG) data produced by the Committee on Best Practice in Data Analysis and Sharing (COBIDAS) (Pernet et al., 2018). The COBIDAS MEEG guidelines were adapted to allow MEEG analysis and reporting to be summarised for the PAF outcome (Supplementary Material 2). Two independent reviewers conducted risk of bias and MEEG methodology assessments. Inconsistencies were resolved by a third reviewer where required.

### 2.9 Data synthesis

Data were synthesised according to the type of NIBS intervention (i.e., separate groups for tACS, rTMS, tDCS). Where interventions had multiple components or conditions with co-interventions (e.g., tDCS with exercise), only the data from the NIBS intervention component were used (McKenzie et al., Cochrane, 2021). When several post-intervention time points were collected (e.g., 5, 10, 15, and 20 minutes post-intervention), the most commonly used time point across studies of the same NIBS intervention was used to avoid issues of multiplicity (McKenzie et al., Cochrane, 2021). When two or more time points were used equally across studies, the earliest time point was used.

#### Meta-analyses

The effect of NIBS interventions on PAF was assessed using mean differences and 95% confidence intervals (CIs) (Borenstein et al., 2009). To synthesise data, random-effects, pair-wise meta-analyses were conducted in RevMan (“Review manager (RevMan) [computer program] version 5.4.” 2000). Weight was given dependent on sample size and amount of variance in outcome within a study. The results of the meta-analyses are presented as forest plots that indicate either increased or decreased PAF following intervention. The distribution of effect sizes were visually examined for each analysis. As studies with small sample sizes were included, and Cohen’s *d* tends to over-estimate standardized mean difference in such cases (Borenstein et al., 2009), *d* was converted to Hedges’ *g* (Hedges, 1981). *The heterogeneity was considered significant when p <* 0.1 in a *χ*^2^ test. *I*^2^ was calculated and values greater than 50% denote important variability across studies that is not due to sampling error (Borenstein et al., 2017, 2009).

#### Subgroup analyses

Based on entrainment principles (Thut et al., 2011a; Vogeti et al., 2022), using stimulation frequencies above, below, or at an individual PAF, should increase, decrease, or have no effect on PAF, respectively. Therefore, tACS conditions were grouped into stimulation frequencies above baseline PAF, below baseline PAF, and those without individualised directions for a subgroup analysis. Due to insufficient numbers of included studies, data were not separated by age, sex, or EEG parameters as planned in the protocol (https://osf.io/2yaxz/).

#### Sensitivity analysis

Sensitivity analyses were not performed by removing studies with high risk of bias (Deeks JJ, Cochrane, 2021) as planned in the protocol (https://osf.io/2yaxz/), because all included studies had low risk of bias.

#### Alternative data synthesis

When meta-analysis was not possible for a particular type of NIBS, an alternative synthesis was conducted based on the Synthesis Without Meta-analysis (SWiM) reporting guidelines (Campbell et al., 2020). For each intervention type, a description of the synthesised findings, the level of certainty in the findings, and possible limitations to the synthesis are described (Campbell et al., 2020).

Where possible, pre- and post-intervention means, SDs mean differences, and standardised mean differences (i.e. estimate of effect), as well as variance and direction of effects on PAF, are reported. Within tables, studies are ordered based on possible differences related to PICO elements (i.e., participant, intervention, control, outcome) to informally highlight possible sources of heterogeneity.

#### Certainty of evidence

Certainty of evidence, as defined in previous studies (Buscemi et al., 2017; Karjalainen et al., 2001; McLean et al., 2010; Scholten-Peeters et al., 2003), forms the basis of prioritising results for summary and conclusion:

- Strong evidence: consistent findings from two or more cohorts with low risk of bias.
- Moderate evidence: consistent findings from at least one cohort with low risk of bias and one or more cohorts with high risk of bias.
- Limited evidence: findings from only one study with low risk of bias or consistent findings in one or more studies with high risk of bias.
- Conflicting evidence: inconsistent findings irrespective of risk of bias.
- No evidence: no studies found.

## 3 Results

### 3.1 Search results

The initial search for all types of interventions used to modulate PAF (e.g., exercise, drugs, NIBS) in EMBASE, PsychINFO, PubMed, Scopus, and the Cochrane library in February 2021 provided a total of 6609 references. After adjusting for duplicates, 4827 abstracts were screened, after which 530 full-texts were screened. A total of 86 studies met the inclusion criteria for all varieties of PAF modulators, nine of which were NIBS interventions. The second search with only NIBS key terms (deviations tracked: https://osf.io/2yaxz/) identified 284 additional references of which 213 abstracts were screened after removing duplicates. Fifty full-texts were screened, resulting in two additional NIBS studies that met the inclusion criteria published between February 2021 and April 2023. Therefore, a total of 11 NIBS studies were identified for inclusion in the review (Anderson et al., 2007; Capotosto et al., 2014; Haberbosch et al., 2019; Kleinert et al., 2017; Okamura et al., 2001; Pahor and Jaušovec, 2016; Ronconi et al., 2020; Sato et al., 2021; Stecher et al., 2021; Stecher and Herrmann, 2018; Steinmann et al., 2022). See Figure 1 for study selection summary.

**Figure 1:**
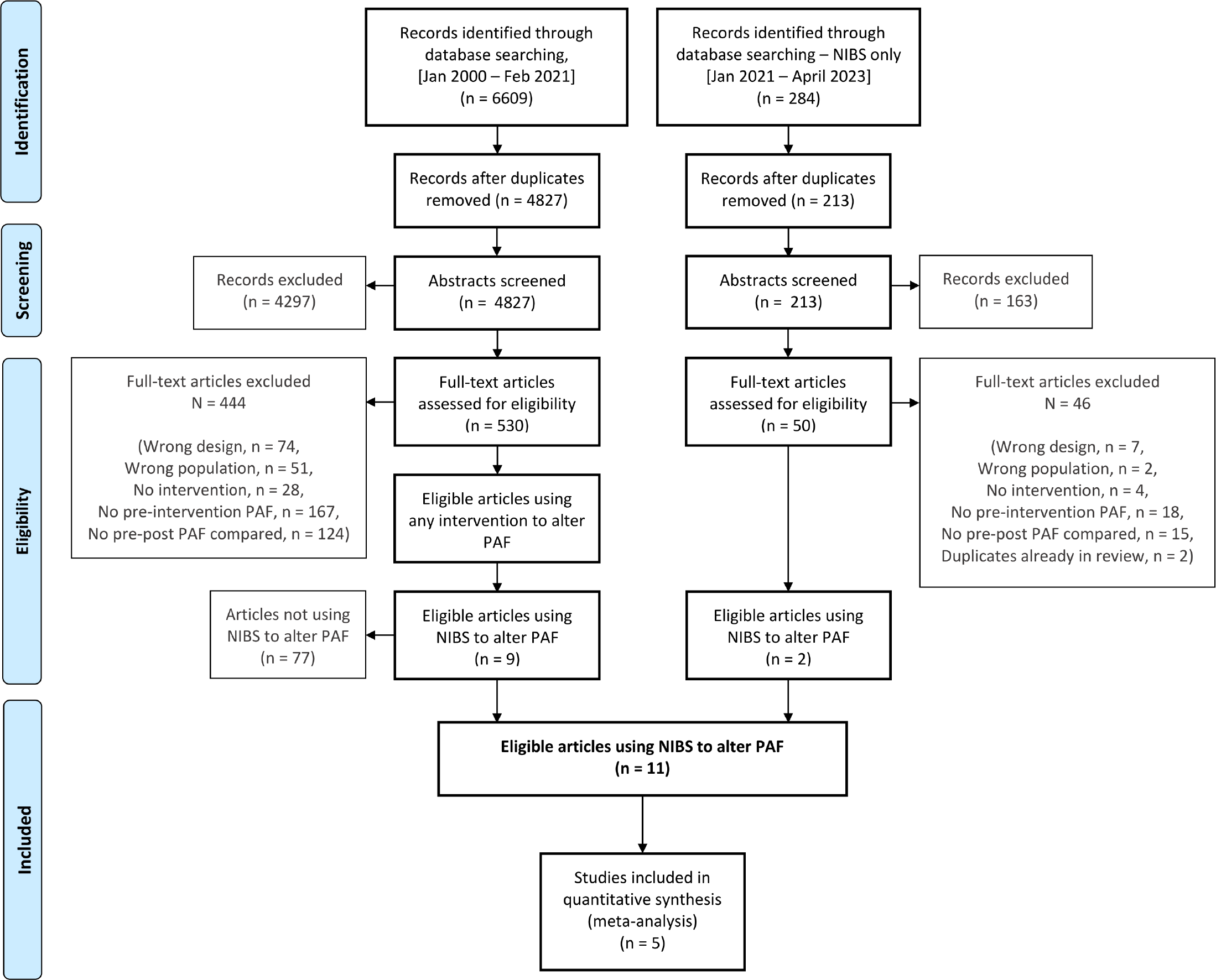
Preferred reporting items for systematic reviews and meta-analysis (PRISMA) flow diagram of the screening and inclusion of studies. *NIBS = non-invasive brain stimulation; PAF = peak alpha frequency*.

In total, the included studies involved 212 healthy adult participants. Participant age was not reported by one study (Anderson et al., 2007). Three types of NIBS interventions were identified: tACS (n = 7 studies) (Haberbosch et al., 2019; Kleinert et al., 2017; Pahor and Jaušovec, 2016; Ronconi et al., 2020; Stecher et al., 2021; Stecher and Herrmann, 2018; Steinmann et al., 2022), rTMS (n = 3 studies) (Anderson et al., 2007; Capotosto et al., 2014; Okamura et al., 2001), and tDCS (n = 1 study) (Sato et al., 2021). The study designs included repeated measures (n = 1), parallel groups (n = 5), and crossover designs (n = 5). Six studies were randomised, controlled, and three studies were non-randomised, controlled. See Table 1 for further details.

**Table 1:**
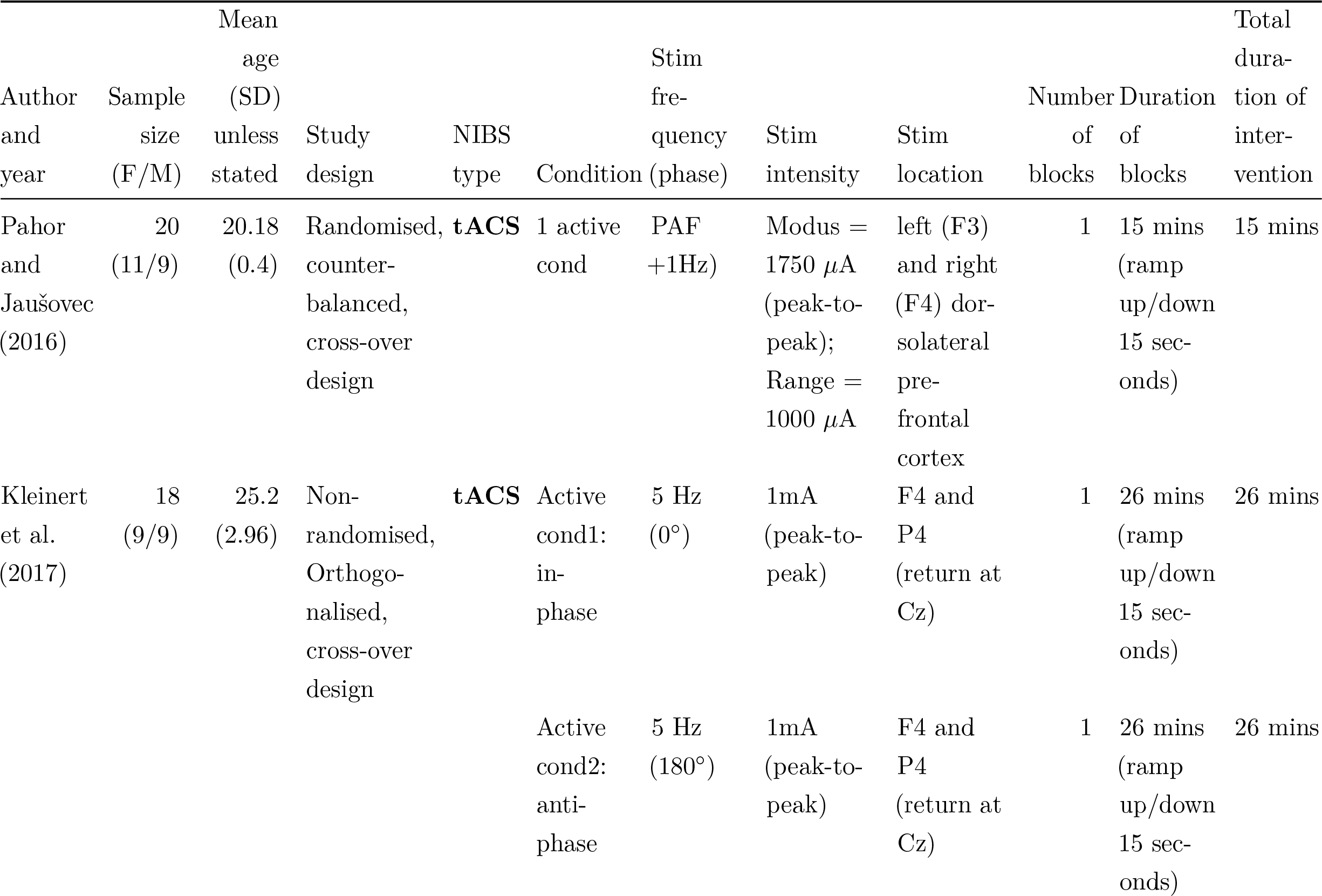
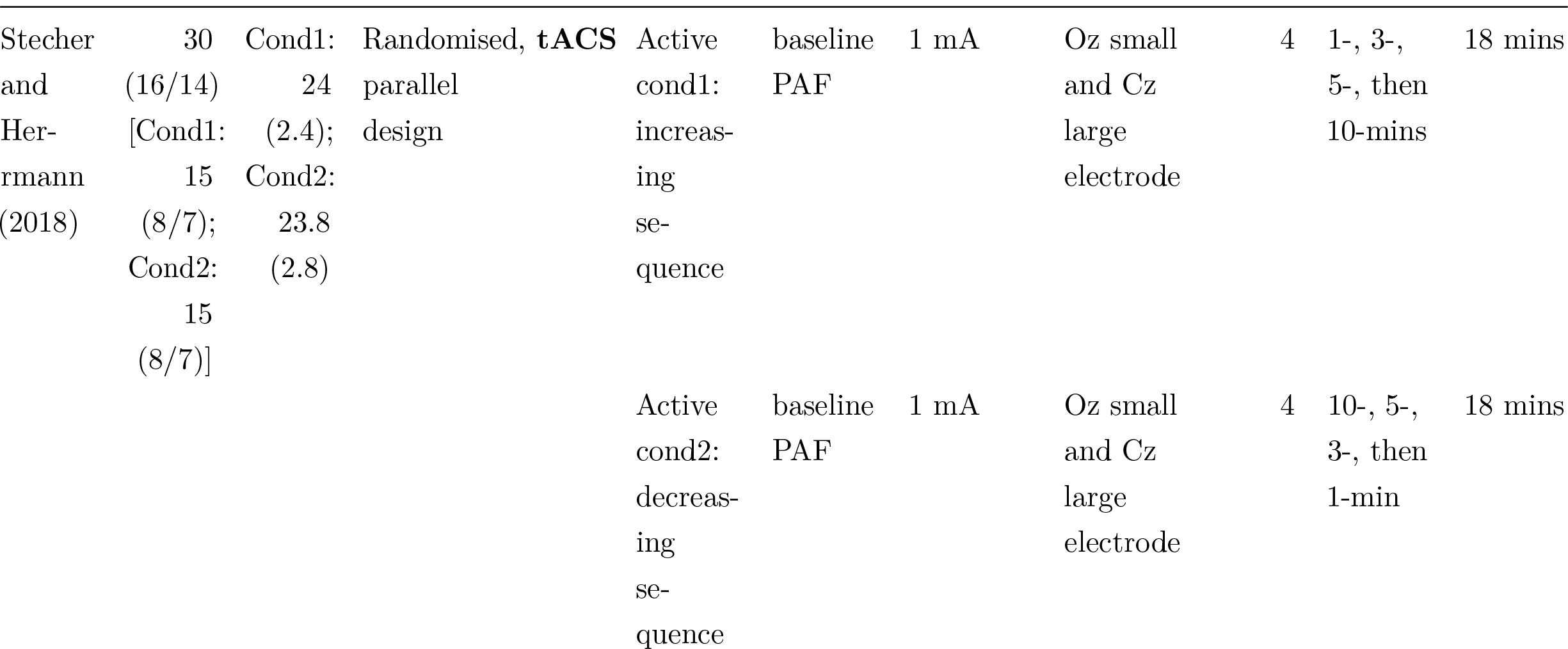
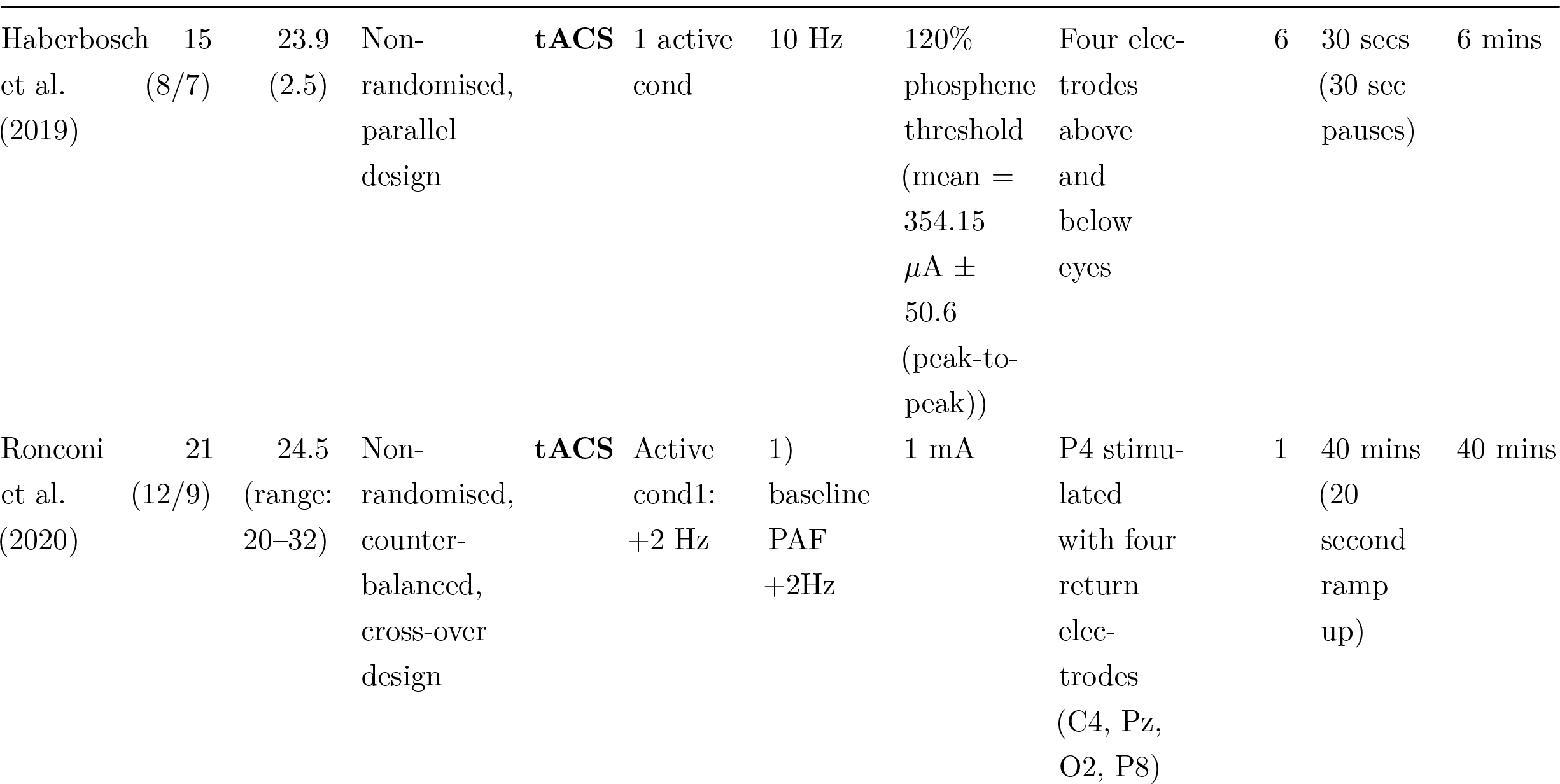
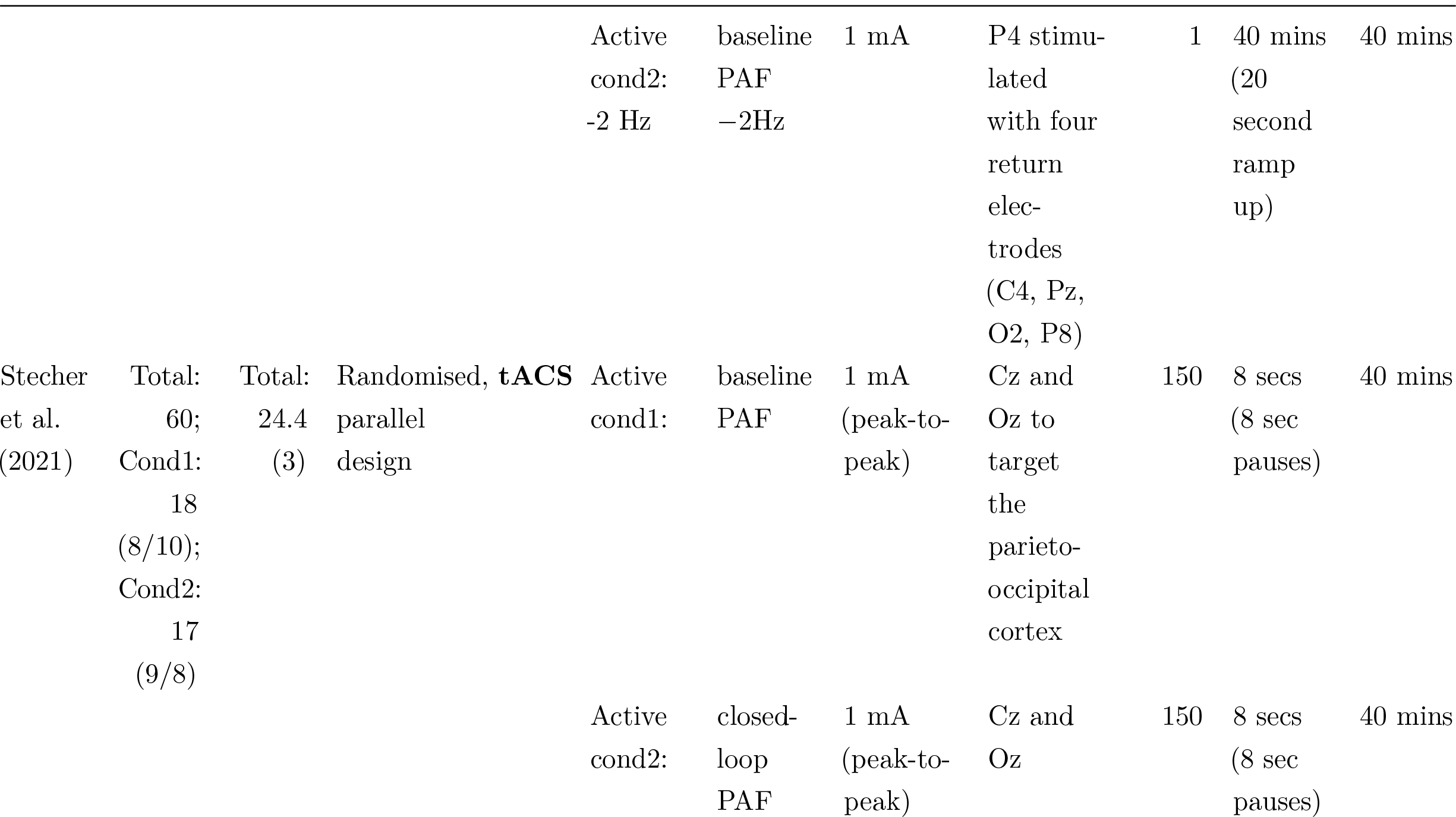
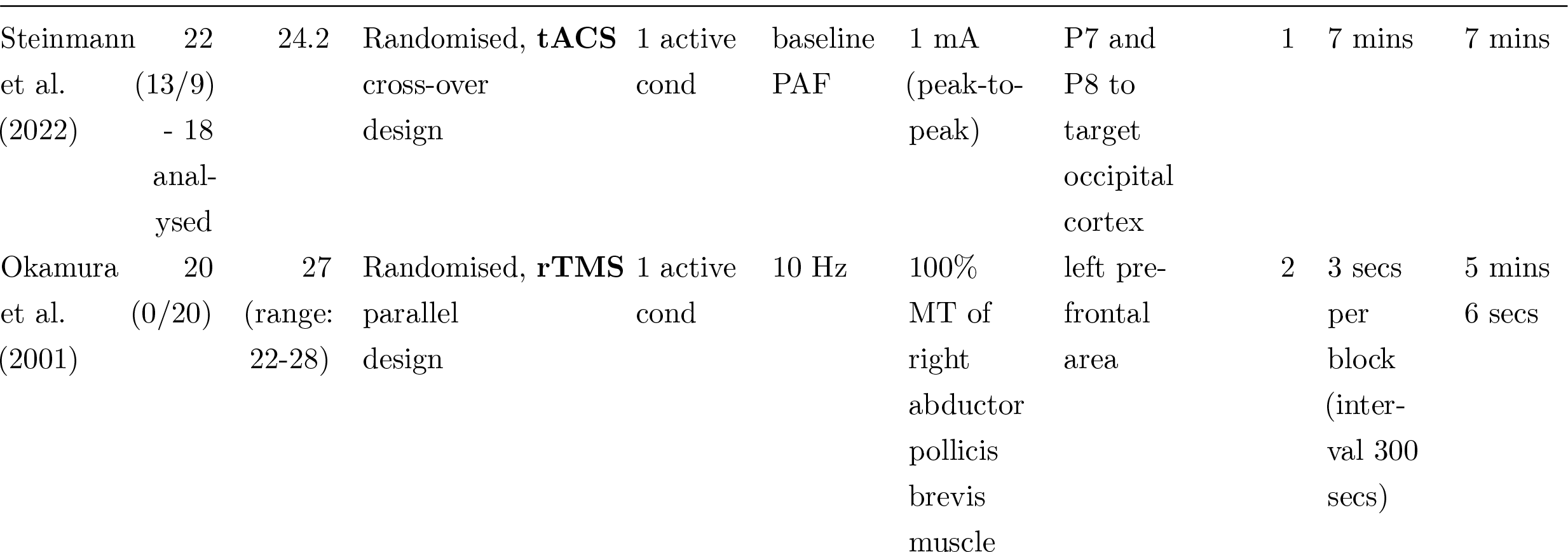
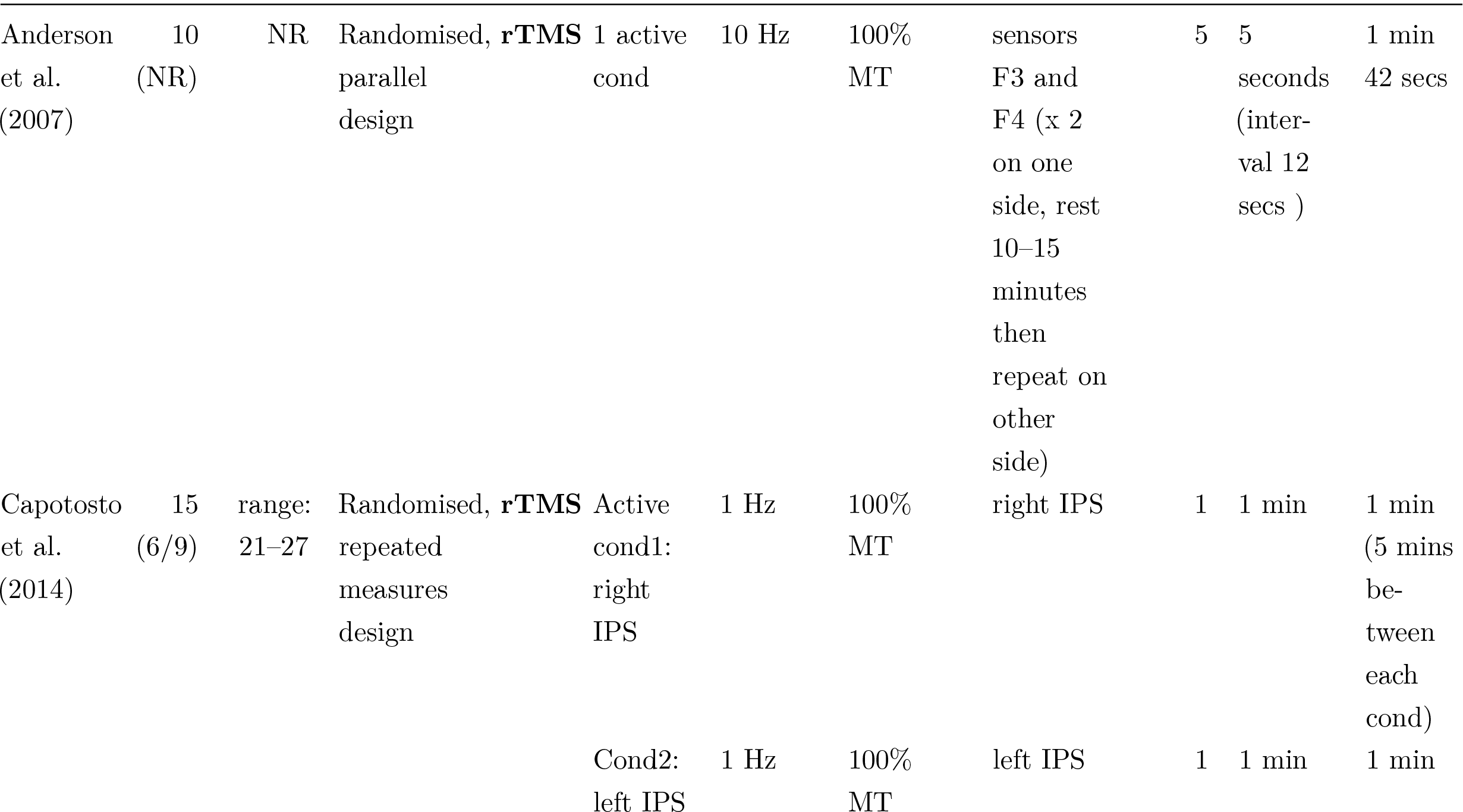
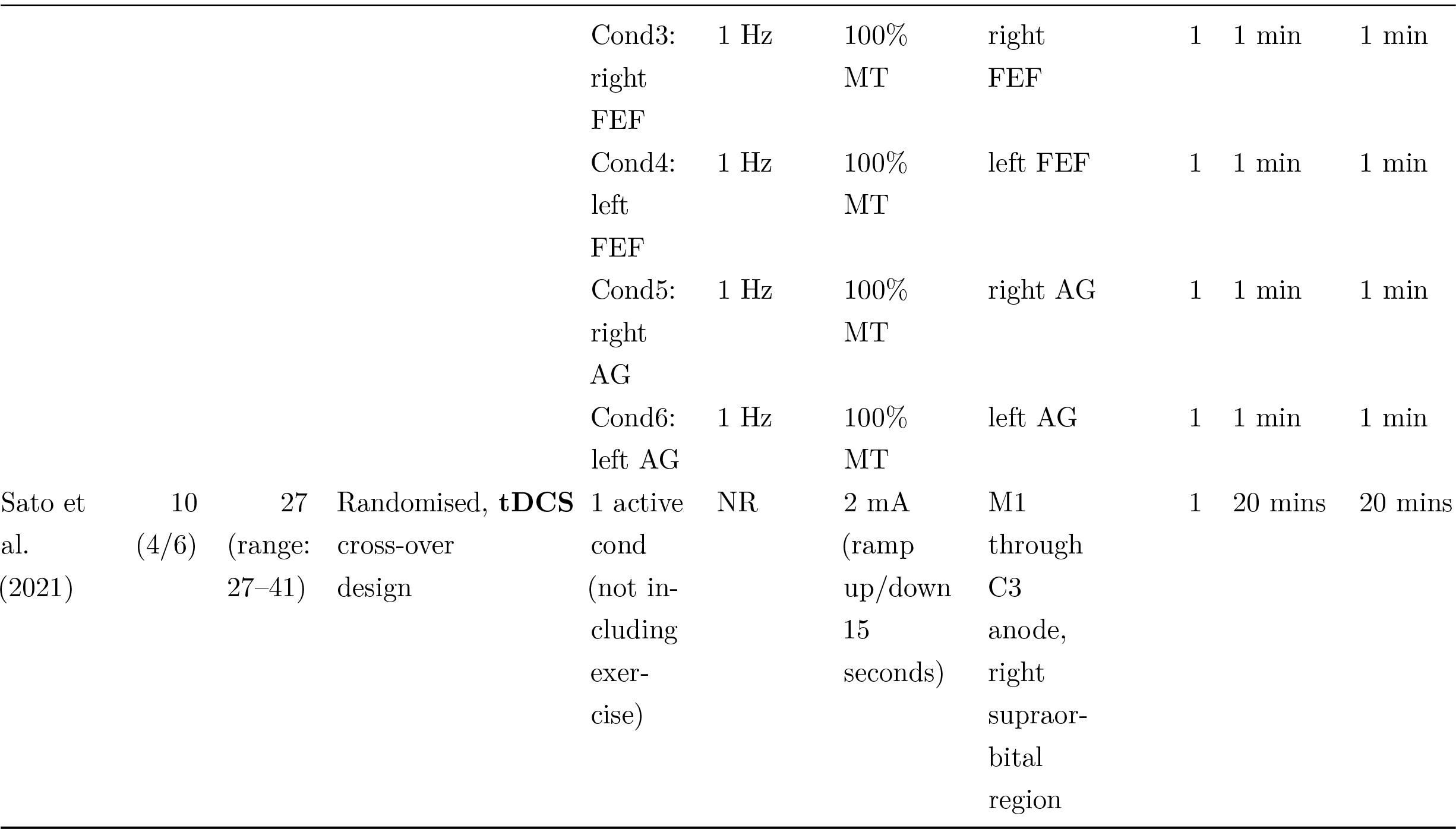
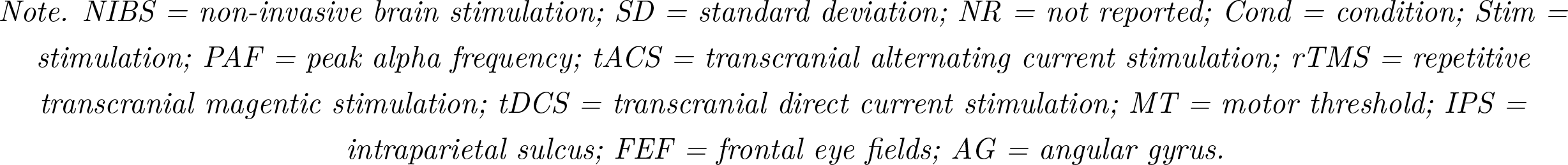
Characteristics of included studies.

### 3.2 EEG methodology and quality

Table 2 shows a summary of EEG methodology. The average score for the EEG methodological assessment was 0.50 out of a maximum score of 1.0 (range: 0.35–0.61). The reliability and precision of PAF measurements can be altered by a variety of factors, such as the recording duration, PAF calculation method, and whether data are recorded with eyes opened or closed (Chiang et al., 2011; Chowdhury et al., 2023; Corcoran et al., 2018).

**Table 2:**
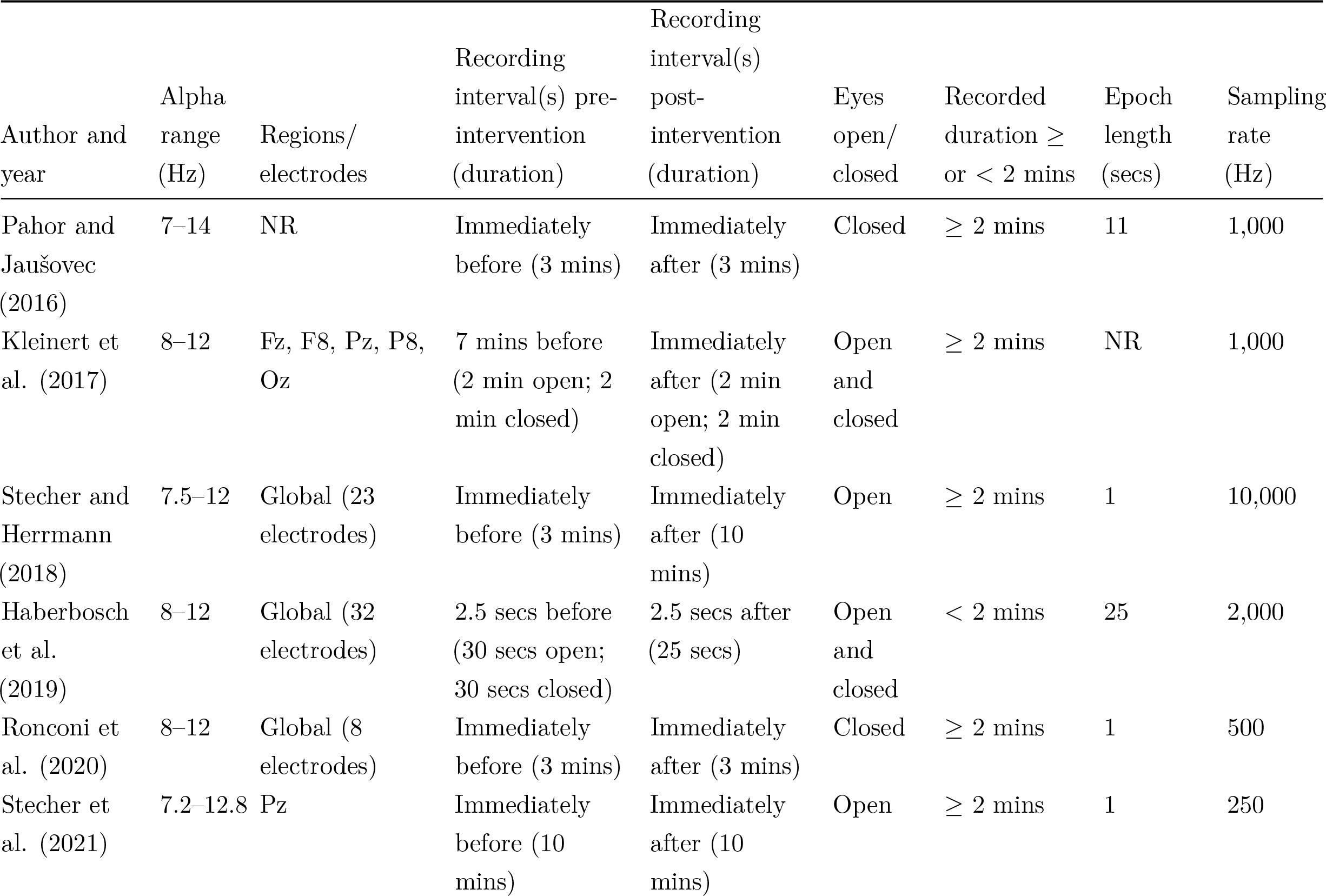
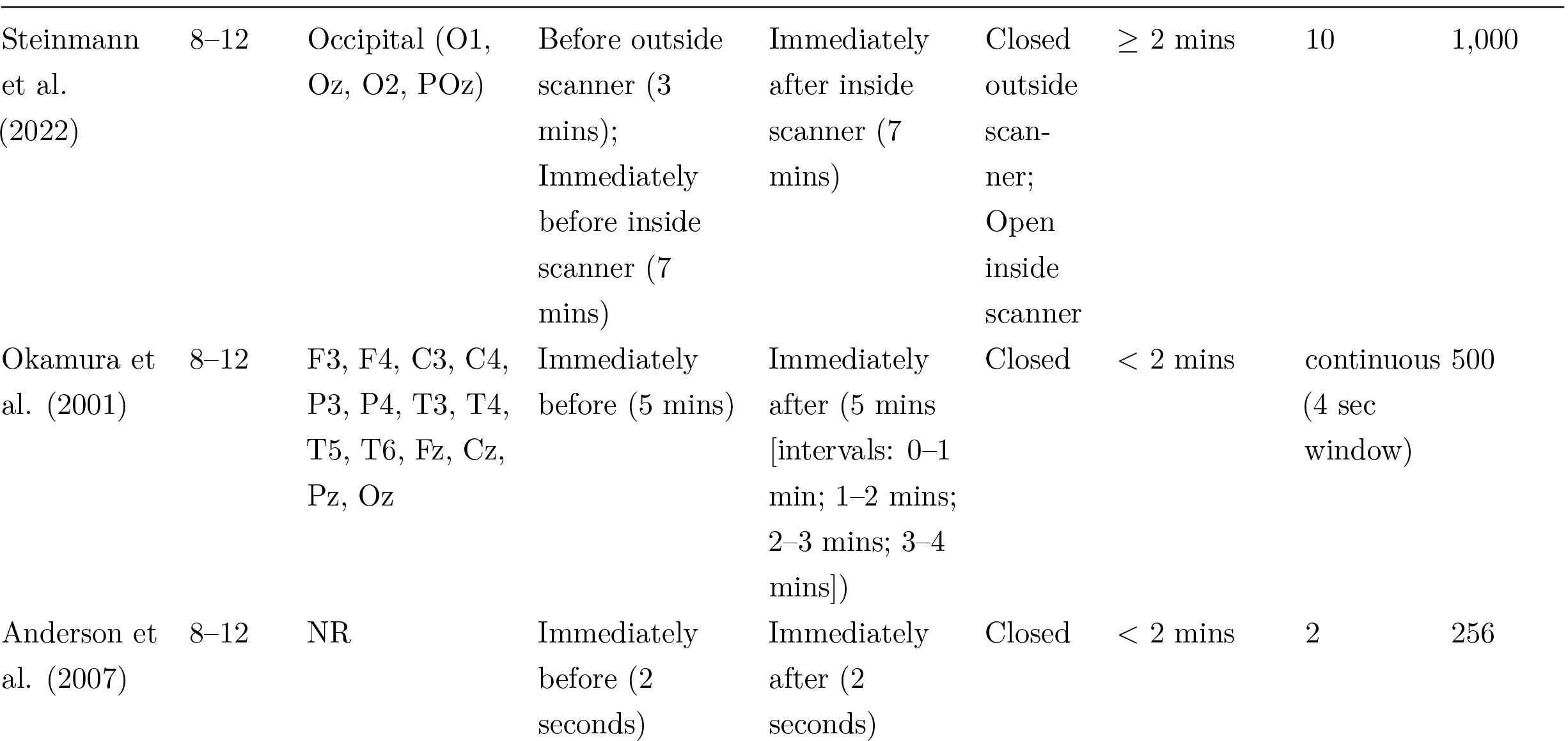
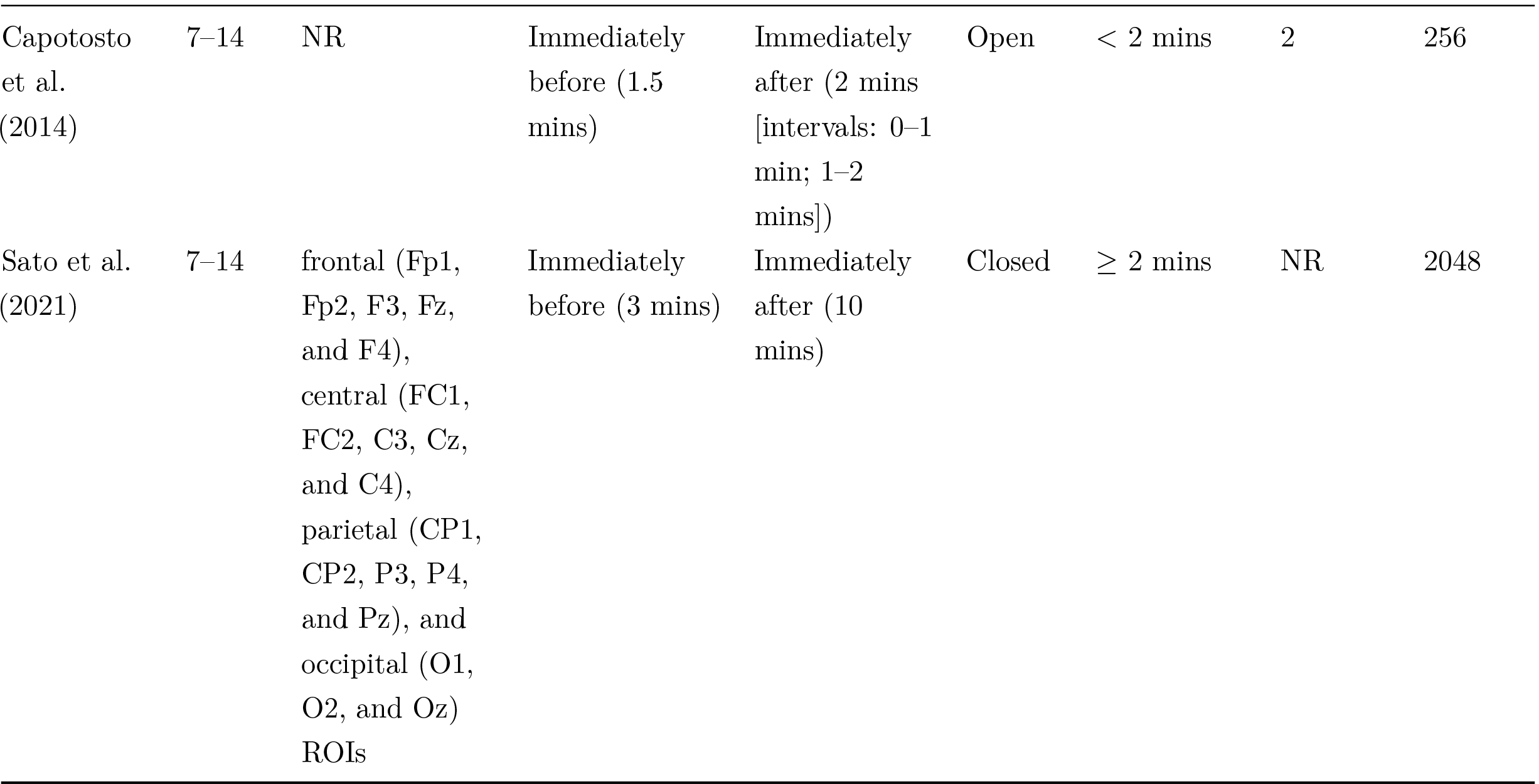
Summary of resting state electroencephalography (EEG) methods.

All studies reported using seated resting state EEG data, recorded immediately before and after NIBS interventions, and all reported whether resting state EEG was recorded with eyes open or eyes closed. Specifically, three studies recorded with eyes open (Capotosto et al., 2014; Stecher et al., 2021; Stecher and Herrmann, 2018), five with eyes closed (Anderson et al., 2007; Okamura et al., 2001; Pahor and Jaušovec, 2016; Ronconi et al., 2020; Sato et al., 2021), and three recorded both conditions (Haberbosch et al., 2019; Kleinert et al., 2017; Steinmann et al., 2022). One study did not have *pure* resting states as participants watched a video during resting states recorded inside an MRI scanner (Steinmann et al., 2022).

Seven studies recorded the resting state EEG for two or more minutes (Kleinert et al., 2017; Pahor and Jaušovec, 2016; Ronconi et al., 2020; Sato et al., 2021; Stecher et al., 2021; Stecher and Herrmann, 2018; Steinmann et al., 2022), while the remainder recorded for less than two minutes (range: 2 seconds–1.5 minutes) (Anderson et al., 2007; Capotosto et al., 2014; Haberbosch et al., 2019; Okamura et al., 2001). No study reported the number of electrodes excluded or interpolated, how many artefact-free epochs were included, or the length of time included per condition for the PAF calculation, except for Haberbosch et al. (2019) who reported 25 seconds of artefact-free data. For PAF calculation, five studies used an 8–12 Hz alpha range (Anderson et al., 2007; Haberbosch et al., 2019; Kleinert et al., 2017; Ronconi et al., 2020; Steinmann et al., 2022), three used 7–14 Hz (Capotosto et al., 2014; Pahor and Jaušovec, 2016; Sato et al., 2021), one used 7.2–12.8 Hz (Stecher et al., 2021), and one used 7.5–12 Hz (Stecher and Herrmann, 2018). The peak picking method of determining PAF was used in all studies. Various electrodes were used for determining PAF (Table 2), with three studies not reporting which electrodes were used (Anderson et al., 2007; Capotosto et al., 2014; Pahor and Jaušovec, 2016).

### 3.3 Risk of bias

One study had ‘some concerns’ regarding overall risk of bias (Steinmann et al., 2022), while all other studies had a ‘low’ risk of bias (Figure 2). Two studies had ‘some concerns’ in the “deviations from the intended interventions” domain, due to a lack of reported information (Haberbosch et al., 2019; Sato et al., 2021). Additionally, two studies had ‘some concerns’ in the “selection of reported results” domain, because they conducted multiple analyses or outcomes without a pre-registered analysis plan (Stecher et al., 2021; Steinmann et al., 2022).

**Figure 2:**
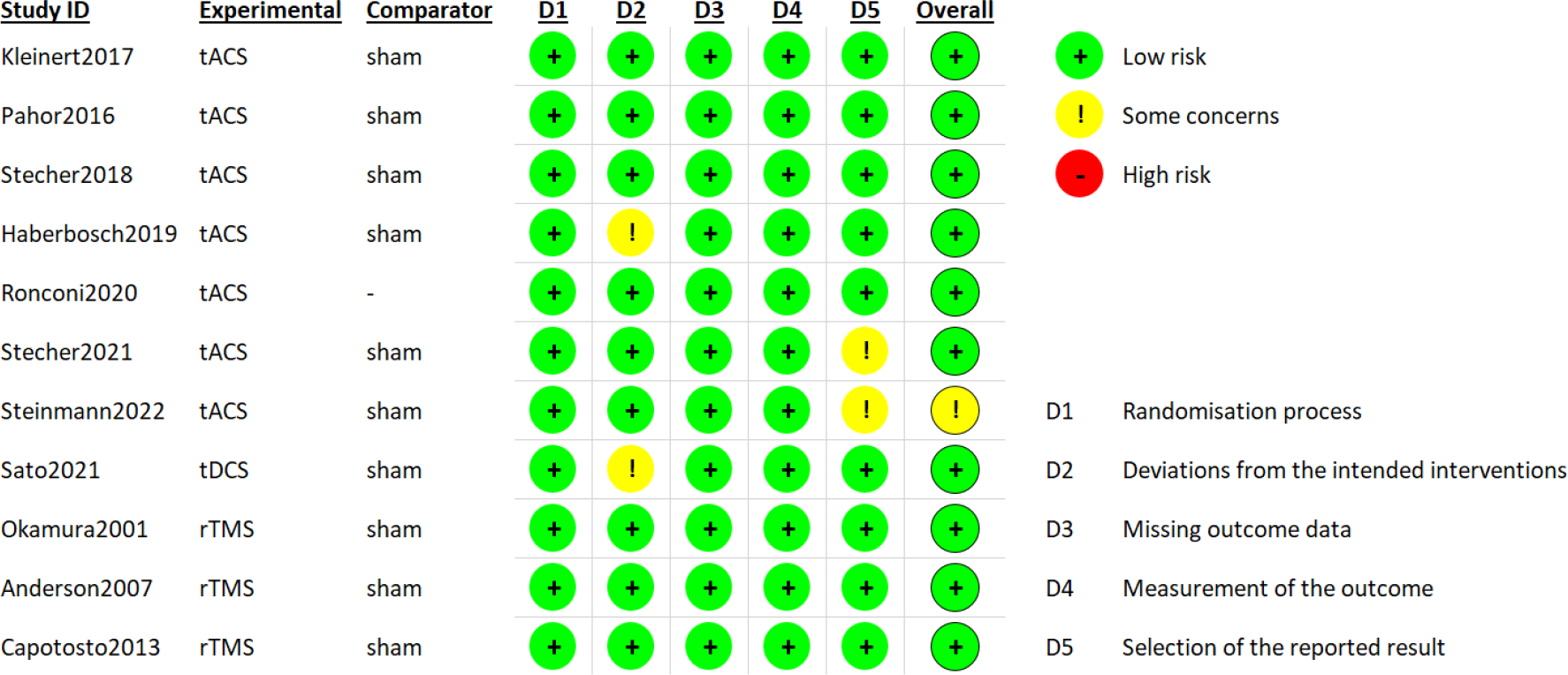
Risk of bias results for included studies, using version 6 of the Cochranes Collaboration’s tool for assessing risk of bias (Higgins et al., 2011; Sterne et al., 2019).

### 3.4 Effects of tACS on PAF

The characteristics of the seven tACS studies are displayed in Table 1. Two authors could not provide pre- and post-tACS PAF means and SDs, so these studies were excluded from the meta-analysis (Kleinert et al., 2017; Steinmann et al., 2022). Pre- and post-tACS PAF means and SDs were collected from eight conditions across the remaining five studies.

#### Meta-analysis of tACS effects on PAF

The eight active tACS conditions included in the meta-analysis varied in stimulation parameters, including stimulation location (i.e. central (Pahor and Jaušovec, 2016; Stecher and Herrmann, 2018), parietal (Ronconi et al., 2020), central-parietal (Stecher et al., 2021), and around the eyes (Haberbosch et al., 2019)), duration (i.e. 6 (Haberbosch et al., 2019), 15 (Pahor and Jaušovec, 2016), 18 (Stecher and Herrmann, 2018), and 40 minutes (Ronconi et al., 2020; Stecher et al., 2021)), and frequency. Stimulation frequencies used included 10 Hz fixed frequency (Haberbosch et al., 2019), above each individual’s PAF (i.e. +1 Hz (Pahor and Jaušovec, 2016) and +2 Hz (Ronconi et al., 2020)), below each individual’s PAF (i.e. -2 Hz (Ronconi et al., 2020)), fixed at individual PAF (Stecher et al., 2021; Stecher and Herrmann, 2018), or a closed-loop PAF stimulation that calculated a new PAF and adjusted stimulation frequency every 8 seconds (Stecher et al., 2021) (Table 1). Meta-analysis showed no change in PAF after tACS intervention (5 studies, 8 conditions, 141 participants, MD = -0.12, 95% CI = -0.32 to 0.08, *Z* = 1.18, *p* = 0.24, *X*^2^(7) = 3.88, *p* = 0.79 *I*^2^ = 0%; Figure 3).

**Figure 3:**
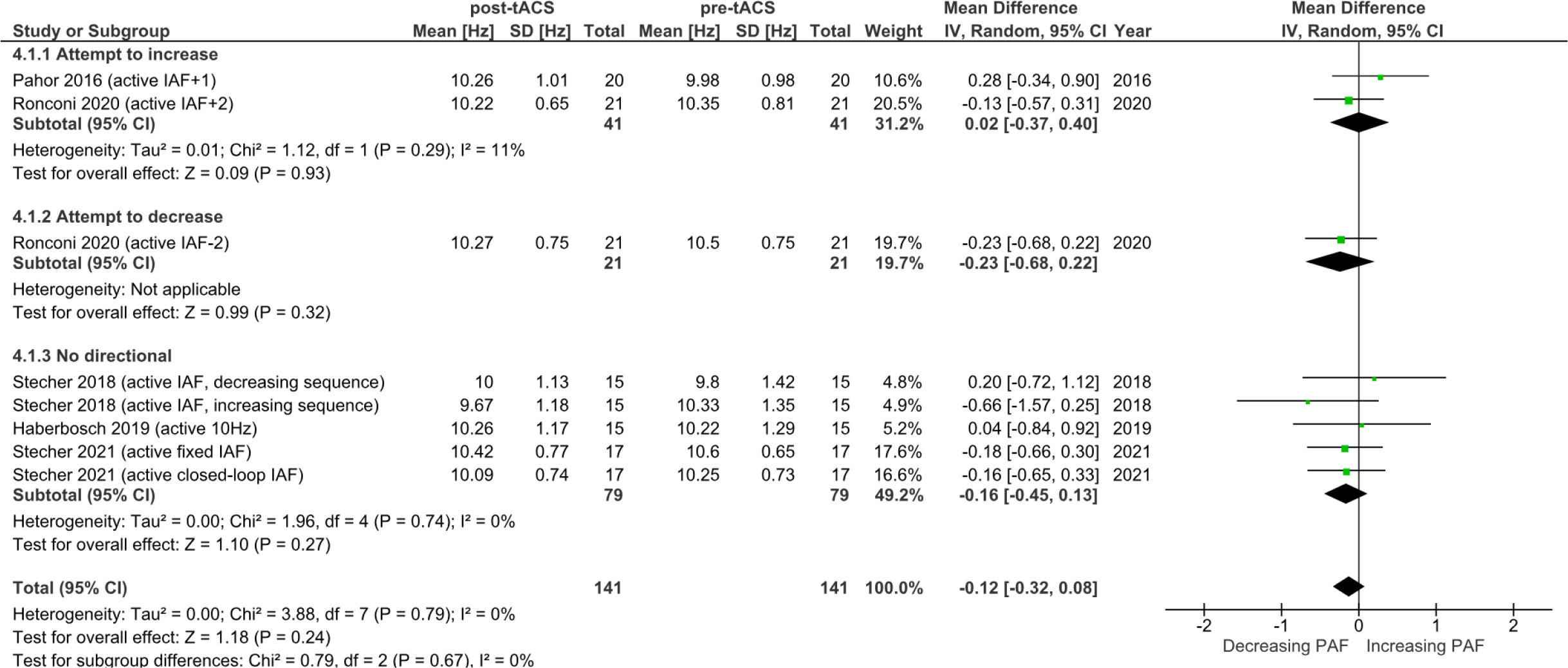
Forest plot showing studies that measured peak/individual alpha frequency (PAF/IAF) pre- and post-transcranial alternating current stimulation (tACS) interventions.

##### Subgroup analysis

Two tACS conditions applied stimulation above individual PAF (i.e. +1 Hz and +2 Hz) (Pahor and Jaušovec, 2016; Ronconi et al., 2020). Pair-wise comparisons showed no increases in PAF (41 participants, MD = 0.02, 95% CI = -0.37 to 0.40, *Z* = 0.09, *p* = 0.93, *X*^2^(1) = 1.12, *p* = 0.29, *I*^2^ = 11%; Figure 3). One condition applied tACS below individual PAF (i.e. -2 Hz) (Ronconi et al., 2020), showing no decreases in PAF (21 participants, MD = -0.23, 95% CI = -0.68 to 0.22, *Z* = 0.99, *p* = 0.32). Five conditions did not individualise tACS (i.e. stimulated at fixed or closed-loop individual PAF (Stecher et al., 2021; Stecher and Herrmann, 2018), or at a fixed 10 Hz (Haberbosch et al., 2019)). No differences in PAF were found after tACS under these conditions (79 participants, MD = -0.16, 95% CI = -0.45 to 0.13, *Z* = 1.10, *p* = 0.27, *X*^2^(4) = 1.96, *p* = 0.74, *I*^2^ = 0%).

### 3.5 Effects of rTMS on PAF

Characteristics of three rTMS studies are displayed in Table 1, (Anderson et al., 2007; Capotosto et al., 2014; Okamura et al., 2001) and are synthesised qualitatively, due to lack of data. Capotosto et al. (2014) applied singular one-minute blocks of 1Hz rTMS at 100% of motor threshold (MT) to six different brain regions separately (i.e. the right and left for intraparietal sulcus, frontal eye fields, and angular gyrus). They reported no evidence for PAF modulation at any location (Capotosto et al., 2014). Anderson et al. (2007) applied five blocks of 10 Hz rTMS, lasting five seconds each block, to the dorsal lateral pre-frontal cortex (DLPFC), finding evidence for an increase in PAF. Okamura et al. (2001) applied two blocks of 10 Hz rTMS, lasting three seconds each block, to the left pre-frontal area, finding evidence for increases in PAF lasting for 1–2 minutes after rTMS. These PAF increases were found in frontal and central sensors (i.e. F3, F4, C3, T3, T4, Fz, and Cz). In summary, there is moderate evidence from two studies that stimulation at 10 Hz to frontal regions could increase PAF (Anderson et al., 2007; Okamura et al., 2001).

### 3.6 Effects of tDCS on PAF

Only one study investigated the effect of tDCS on PAF with 10 participants (Sato et al., 2021) (Table 1). Sato et al. (2021) applied 2mA anodal tDCS for a singular 20-minute block to the left M1, finding no effect on PAF in any of the four regions of interest (i.e., frontal, central, parietal, occipital).

## 4 Discussion

This is the first systematic review to synthesise the evidence for the effect of NIBS interventions on PAF speed. Eleven studies using tACS (seven studies), rTMS (three studies), and tDCS (one study) were included. There was moderate evidence for increased PAF speed from two studies applying rTMS at 10 Hz to frontal regions and no evidence that tACS or tDCS modulated PAF speed. Overall, the evidence is limited, and heterogeneity of stimulation parameters hampers conclusions.

### 4.1 tACS does not alter PAF

tACS delivers sinusoidal currents to the brain at a specific frequency and is thought to modify synaptic activity by rhythmically altering neuron membrane potentials and the likelihood of neuronal firing, offering the potential to modulate brain oscillations (Bergmann and Hartwigsen, 2021; Frohlich and Riddle, 2021; Fröhlich and McCormick, 2010; Herrmann et al., 2013; Thut et al., 2011a; Vogeti et al., 2022). However, there was no evidence of an effect of tACS on PAF in this review (Haberbosch et al., 2019; Kleinert et al., 2017; Pahor and Jaušovec, 2016; Ronconi et al., 2020; Stecher et al., 2021; Stecher and Herrmann, 2018; Steinmann et al., 2022). Definitive conclusions are limited due to the scarcity of studies, small sample sizes, and substantial heterogeneity in stimulation parameters, including frequency (i.e. at individual fixed or closed-loop PAF, PAF +2 Hz, PAF +1 Hz, PAF –2 Hz, fixed 10 Hz, fixed 5 Hz), duration (range: 6–40 minutes), and location (i.e. frontal, parietal, occipital). Given that parameter configurations are likely to directly influence modulation of oscillations (Polanía et al., 2018; Thut et al., 2011b; Vogeti et al., 2022), future research should systematically investigate the effects of these parameters on PAF.

Entrainment describes the frequency and phase alignment between oscillatory systems (Pikovsky et al., 2001). In the context of brain oscillations, entrainment refers to the process by which the frequency and phase of neural oscillations synchronise to an external rhythmic stimulus (Frohlich and Riddle, 2021; Lakatos et al., 2019; Pikovsky et al., 2001; Thut et al., 2011a). Based on the theory of entrainment, stimulation frequencies faster than an individual’s PAF should increase PAF speed, while slower frequencies should decrease it (Lakatos et al., 2019; Thut et al., 2011b). Subgroup analysis found no effect on PAF, regardless of the stimulation frequency used. Notably however, the largest subgroup stimulated either at a non-individualised frequency (i.e. 10 Hz) or at individual PAF (e.g. fixed or closed-loop). As entrainment theory would predict no change in PAF under these conditions, the insignificant finding is perhaps unsurprising. Only one condition stimulated below individual PAF (Ronconi et al., 2020), while two conditions stimulated above (Pahor and Jaušovec, 2016; Ronconi et al., 2020), also showing no change in PAF. A recent study corroborated our lack of findings, reporting no effect of left posterior parietal tACS on PAF (N = 21) across various stimulation frequencies (i.e. PAF, PAF +2 Hz, PAF –2 Hz, and sham stimulation) in a randomised cross-over design (Kemmerer et al., 2022). This study was excluded from the current review due to inclusion of participants older than 65 years (see Materials and methods).

It is plausible that the lack of overall effect stems from the application of stimulation frequencies too far from an individual’s baseline PAF, such as PAF *±*2 Hz (Kemmerer et al., 2022; Ronconi et al., 2020). According to the physical principles of synchronisation (Pikovsky et al., 2001), it is easier to entrain internal rhythms with external rhythms that closely match in frequency (Thut et al., 2011a; Vogeti et al., 2022); with existing evidence that higher stimulation intensities are required to achieve entrainment when the frequency is not closely matched (Ali et al., 2013; Frohlich and Riddle, 2021; Negahbani et al., 2019). However, a comprehensive investigation into the effects of stimulation frequencies above and below individual PAF on PAF modulation is lacking. Future tACS research should use larger sample sizes and adopt smaller increments of stimulation frequencies around individual baseline PAF, such as PAF *±*0.2 Hz or PAF *±*0.5 Hz, to provide more detailed understanding of the effects of tACS stimulation frequency on PAF.

### 4.2 rTMS at 10 Hz may increase PAF

Two studies used 10 Hz rTMS over pre-frontal regions, finding transient increases in PAF (∼1.5 Hz) in frontal, central, and temporal EEG electrodes lasting for 2 minutes (Anderson et al., 2007; Okamura et al., 2001), suggesting localised effects near the stimulation site. Conversely, one study that examined 5 Hz rTMS over multiple locations found no evidence for PAF modulation (Capotosto et al., 2014). However, caution is needed when interpreting these results due to small sample sizes and the limited number of studies. As well as replication studies, future research could use longer stimulation durations or multiple sessions in larger samples to assess whether sustained change in PAF can be induced by rTMS.

Included rTMS studies employed fixed 5 or 10 Hz frequencies; however, literature on PAF modulation during stimulation suggests more robust effects may be induced by applying individualised rTMS frequencies (i.e. PAF *±*1 Hz), as indicated by Di Gregorio et al. (2022). Furthermore, studies on rTMS in depression suggest that the proximity of an individual’s PAF to the external driving rhythm (i.e. rTMS frequency) has a quadratic relationship with improvements in depression symptoms (Corlier et al., 2019; Roelofs et al., 2021), such that patients with baseline PAF closer to the stimulation frequency of 10 Hz had greater improvements in depressive symptoms than those with baseline PAF further from 10 Hz. This indicates that the relationship between baseline PAF and stimulation frequency may impact functional outcomes related to PAF speed, alongside the magnitude and direction of PAF change (Corlier et al., 2019; Roelofs et al., 2021). However, these studies did not assess whether PAF was modulated by the stimulation. Future research should investigate effects of individualised stimulation frequencies on PAF and the relationship between baseline PAF and stimulation frequency of rTMS.

### 4.3 tDCS does not alter PAF

Research investigating the effect of tDCS interventions on PAF is scarce. One study (Sato et al., 2021) examined the effect of 2 mA tDCS over the motor cortex on PAF in 10 participants, reporting no effect on PAF. The authors noted that future investigations should reduce the time between stimulation with tDCS and measurement with EEG to enhance the assessment of tDCS effects on PAF (Sato et al., 2021).

### 4.4 Recommendations and future directions

Based on this review, future research should: 1) explore smaller increments of stimulation frequencies for tACS around individuals’ PAF, such as PAF *±*0.2 Hz, or PAF *±*0.5 Hz; 2) explore the relationship between baseline PAF and stimulation frequency for tACS and rTMS; 3) use longer durations or multiple sessions of 10 Hz rTMS; and 4) explore individualised stimulation frequencies for rTMS.

In addition, future research should consider the theoretical assumptions underlying NIBS mechanisms on brain oscillations, specifically, the theory of spike-timing dependent plasticity (STDP) alongside entrainment (Vogeti et al., 2022). While entrainment refers to synchronisation of endogenous oscillations to an external driving frequency, STDP suggests that stimulation leads to synaptic changes based on the timing of neuronal firing in the targeted region (see Vogeti et al. (2022) for a full review). The choice of stimulation parameters, such as the stimulation intensity or timing and duration of stimulation trains, may have varying effects on PAF depending on which theory or combination of theories are employed (Bergmann and Hartwigsen, 2021; Vogeti et al., 2022). While the mechanism through which NIBS may impact PAF remains uncertain, future research should carefully consider the impact of these theories on the selection of stimulation parameters.

Lastly, future studies should consider the influence of EEG recording, pre-processing, and PAF calculation methods. In the current review, included studies employed varying locations, durations, and resting state conditions (i.e. eyes closed or eyes open) for EEG measurements. These discrepancies in EEG methodology have the potential to influence PAF values and study outcomes (Chowdhury et al., 2023; Corcoran et al., 2018; Furman et al., 2021; Gil Avila et al., 2023; McLain et al., 2022). For example, recording resting state EEG with eyes closed emphasises the contribution of occipital alpha oscillations, as opposed to when eyes are open (Berger, 1929). In addition, the peak picking method of PAF estimation in the included studies is less reliable than the centre of gravity (CoG) method (Chowdhury et al., 2023) and also overlooks the possibility of multiple alpha peaks in an individual’s frequency spectra (Chiang et al., 2011, 2008). This oversight disregards the potential for changes in PAF speed through relative increases in the power of faster alpha oscillations compared to slower alpha oscillations (Furman et al., 2021). Future studies should carefully consider the conceptualisation, measurement, and quantification of PAF when evaluating the effect of NIBS interventions on PAF. Moreover, it is imperative for future work to adhere to published reporting guidelines (Pernet et al., 2020, 2018), consider data sharing (Pernet et al., 2020; Ploner et al., 2017), and adopt standardised (Gil Avila et al., 2023) or no-clean (Chowdhury et al., 2023) pre-processing pipelines to enhance research quality, transparency (Cohen, 2017b), and facilitate comparison of PAF values across studies (McLain et al., 2022).

### 4.5 Study limitations and constraints

There are several limitations in this review. First, the included studies had small sample sizes and likely had low statistical power. Second, the generalisability of the results to patient populations cannot be inferred, as studies were restricted to healthy populations. Third, the review only considered articles published since 2000, potentially missing relevant work published prior to this period. Lastly, the review did not explore the influence of alpha power, despite the extensive literature on NIBS effects on alpha power (Vogeti et al., 2022). Investigating the interactions between power and alpha peaks after NIBS could provide valuable insights into PAF modulation.

### 4.6 Conclusion

This systematic review provides preliminary evidence for transient increases in PAF speed after one session of 10 Hz rTMS. Further investigations are warranted to assess the sustained modulation of PAF by 10 Hz rTMS, using multiple sessions or extended stimulation durations. Although tACS did not influence PAF in this analysis, the mechanism of action makes it theoretically likely to influence PAF speed and further studies are warranted. Exploring variations in stimulation parameters (e.g., frequency, intensity, or duration) within tACS interventions could uncover its capacity to affect PAF. Future studies should delve into optimal protocols and parameter settings for rTMS and tACS, while accounting for individual differences. This research has the potential to not only advance our understanding of PAF modulation through NIBS but also to refine existing therapeutic NIBS interventions for conditions associated with slower PAF, such as depression and chronic pain.

## Notes

### Competing Interest Statement

The authors have declared no competing interest.

