## Supplementary Material 1 for "Can non-invasive brain stimulation modulate peak alpha frequency in the human brain? A systematic review and meta-analysis"

PAF modulation systematic review - database searches

Search strategies for Embase (OVID), PsychINFO (ProQuest), Pubmed (NIH, National Library of Medicine), Scopus (Elsevier), The Cochrane Library (Cochrane Collaboration), and grey literature. All databases can be accessed through UNSW's list of databases.

**EMBASE** (Ovid)

1. (((peak$1 or alpha or alpha-peak or peak-alpha or dominant or individual or mu) adj2 (frequenc* or rhythm*)) or ((dominant or alpha or mu) adj2 peak$1) or (PAF or IAF or IAPF or main frequenc* or mean peak or individual peak)).ab,ti,kw.
2. exp electroencephalogram/ or exp electroencephalography/ or EEG.mp. or electroencephalogra*.mp. or electro-encephalogra*.mp.
3. exp magnetoencephalography/ or (MEG or magnetoencephalogra* or magneto-encephalogra*).mp.
4. 2 or 3
5. ((brain* or cortex* or cortical* or crani* or transcrani* or magneti* or burst or non-invasive or noninvasive) adj4 (stimulat* or electrostim* or electro-stim* or electrotherap* or electro-therap*)).mp.
6. (TMS or rTMS or tPEMF or LFMS or TBS or iTBS or cTBS).mp.
7. (tDCS or tACS or tPCS or tRNS or ECT or CES).mp.
8. 5 or 6 or 7
9. (modulat* or therap* or treatment* or intervention*).mp.
10. (neuro-feedback* or neurofeedback* or bio-feedback* or biofeedback*).mp.
11. exp physical activity/ (relax* or meditat* or exercis*).mp.
12. exp sensory stimulation/ or ((auditory or visual* or nocicept* or somatosensor* or tactil* or light* or photic*) adj3 (stim* or flicker)).tw. or (music* or task$1).mp. or
13. 9 or 10 or 11 or 12
14. exp "agents interacting with transmitter, hormone or drug receptors"/
15. exp central nervous system agents/
16. exp "general and inorganic chemicals"/
17. exp organic compound/
18. exp pharmacology/ or (pharma* or drug*).mp.
19. exp nicotine/ or exp "smoking and smoking related phenomena"/ or exp alcohol/ or (nicotin* or smoking* or alcohol*).mp.
20. 14 or 15 or 16 or 17 or 18 or 19
21. 8 or 13 or 20
22. 1 and 4 and 21
23. limit 22 to english language
24. limit 24 to "humans only (removes records about animals)"

**PsycINFO** (ProQuest)

ab,ti,if("peak frequency" OR "alpha peak" OR "peak alpha" OR "dominant frequency" OR "dominant rhythm" OR "dominant peak" OR "individual peak" OR "mu frequency" OR "PAF" OR "iPAF" OR "iAPF")

AND

(MAINSUBJECT.EXACT("Electroencephalography") OR MAINSUBJECT.EXACT.EXPLODE("Electroencephalography") OR MAINSUBJECT.EXACT("Magnetoencephalography") OR MAINSUBJECT.EXACT.EXPLODE("Magnetoencephalography") OR ab,ti,if((EEG OR electroencephalogra* OR MEG OR magenetoencephalogra*)))

AND

((MAINSUBJECT.EXACT.EXPLODE("Treatment") OR MAINSUBJECT.EXACT.EXPLODE("Stimulation") OR MAINSUBJECT.EXACT.EXPLODE("Feedback") OR MAINSUBJECT.EXACT.EXPLODE("Arts") OR MAINSUBJECT.EXACT.EXPLODE("Behavior") OR MAINSUBJECT.EXACT.EXPLODE("Physical Activity") OR MAINSUBJECT.EXACT.EXPLODE("Drugs") OR MAINSUBJECT.EXACT.EXPLODE("Chemicals") OR MAINSUBJECT.EXACT.EXPLODE("Pharmacology") OR MAINSUBJECT.EXACT.EXPLODE("Nutrition") OR ab,ti,if(TMS OR rTMS OR tPEMF OR LFMS OR TBS OR iTBS OR cTBS OR tDCS OR tACS OR tPCS OR tRNS OR ECT OR CES OR neurofeedback OR "neuro feedback" OR biofeedback OR "bio feedback" OR meditat* OR music* OR danc* OR relax* OR exercise* OR task OR drug* OR pharma* OR smoking* OR nicotin* OR alcohol*)) OR (ab,ti,if(brain* OR cortex* OR cortical* OR crani* OR transcrani* OR magneti*) NEAR/2 ab,ti,if(stimulat* OR electrostim* OR "electro stimulation" OR electrotherap* OR "electro therap*")) OR (ab,ti,if(auditory OR visual* OR nocicept* OR sensory OR somatosensor* OR tactil* OR light* OR photic*) NEAR/2 ab,ti,if(stim* OR flicker)))

Limit to English language

Limit to human population

Limit to adulthood (18 yrs & older) age

**PubMed** (NIH, National Library of Medicine)

Title /Abstract search with MeSH terms

#1) Peak alpha frequency[Title/Abstract] OR peak frequency[Title/Abstract] OR alpha peak[Title/Abstract] OR peak alpha[Title/Abstract] OR dominant frequency[Title/Abstract] OR dominant rhythm[Title/Abstract] OR dominant peak[Title/Abstract] OR individual peak[Title/Abstract] OR mu frequency[Title/Abstract] OR PAF[Title/Abstract] OR iPAF[Title/Abstract] OR iAPF[Title/Abstract]

#2) electroencephalography[Title/Abstract] OR EEG[Title/Abstract] OR magnetoencephalography[Title/Abstract] OR MEG[Title/Abstract] OR electroencephalography[MeSH Terms] OR magnetoencephalography[MeSH Terms]

#3) therapeutics[MeSH Terms] OR therapeutic*[Title/Abstract] OR brain stimulation[Title/Abstract] OR tACS[Title/Abstract] OR tDCS[Title/Abstract] OR TMS[Title/Abstract] OR rTMS[Title/Abstract] OR ECT[Title/Abstract] OR LFMS[Title/Abstract] OR tPEMF[Title/Abstract] OR TBS[Title/Abstract] OR iTBS[Title/Abstract] OR cTBS[Title/Abstract] OR tPCS[Title/Abstract] OR tRNS[Title/Abstract] OR CES[Title/Abstract] OR sensory stimulation[Title/Abstract] OR modulat*[Title/Abstract] OR alter*[Title/Abstract] OR psychiatric somatic therapies[MeSH Terms] OR pharmaceutical preparations[MeSH Terms] OR heterocyclic compounds[MeSH Terms] OR nicotin*[Title/Abstract] OR smoking[Title/Abstract] OR organic chemicals[MeSH Terms] OR alcohol*[Title/Abstract] OR pharmacology[MeSH Subheading] OR psychotherapy[MeSH Terms] OR drug*[Title/Abstract] OR pharma*[Title/Abstract] OR music*[Title/Abstract] OR neurofeedback[Title/Abstract] OR “neuro feedback”[Title/Abstract] OR biofeedback[Title/Abstract] OR “bio feedback”[Title/Abstract] OR meditat*[Title/Abstract] OR exercis*[Title/Abstract] OR physical activit*[Title/Abstract] OR relax*[Title/Abstract] OR behavior and behavior mechanisms[MeSH Terms] OR smoking[Title/Abstract] OR task[Title/Abstract]

#4) #1 AND #2 AND #3

#5) (animals[MeSH Terms]) NOT (humans[MeSH Terms])

#5) #4 NOT #5

Filters: **English** Sort by: **Most Recent**

Don’t apply other limits, because new publications will not have MEDLINE indexing yet.

**Scopus** (Elsevier)

1. ((TITLE-ABS-KEY((peak or alpha-peak or peak-alpha or dominant or individual or mu) W/1 (frequenc* or rhythm*)) OR TITLE-ABS-KEY((dominant or alpha or mu) W/1 peak)OR TITLE-ABS-KEY(PAF or iAF or iAPF or iPAF)))

AND

2. ((TITLE-ABS-KEY(electroencephalogra* or "electro encephalogra*" or EEG) OR TITLE-ABS-KEY(magnetoencephalogra* or "magneto encephalogra*" or MEG)OR TITLE-ABS-KEY(MEEG)))

AND

3. (((TITLE-ABS-KEY((brain* or cortex* or cortical* or crani* or transcrani* or magneti* or burst or "non invasive" or noninvasive) W/3 (stimulat* or electrostim* or "electro stim*" or electrotherap* or "electro therap*")) OR TITLE-ABS-KEY(TMS or rTMS or tPEMF or LFMS or TBS or iTBS or cTBS)OR TITLE-ABS-KEY(tDCS or tACS or tPCS or tRNS or ECT or CES))) OR ((TITLE-ABS-KEY(modulat* or therap* or treatment or intervention) OR TITLE-ABS-KEY(relax* or meditat* or exercis* or music* or task)OR TITLE-ABS-KEY((auditory or visual* or nocicept* or somatosensor* or tactil* or light* or photic*) W/2 (stim* or flicker)))) OR ((TITLE-ABS-KEY(chemical* or pharma* or drug or nicotin* or smoking or alcohol) OR TITLE-ABS-KEY("neuro feedback*" or neurofeedback* or "bio feedback*" or biofeedback*))))

Limit to English language

**Cochrane Central Register of Controlled Trials (CENTRAL)** (Cochrane Collaboration)

1. ((((peak or alpha-peak or peak-alpha or dominant or individual or mu) NEAR (frequenc* OR rhythm*)) OR ((dominant OR alpha OR mu) NEAR peak) OR PAF OR iAF OR iAPF OR iPAF)):ti,ab,kw
2. ((EEG OR electroencephalogra* OR MEG OR magnetoencephalogra* OR MEEG)):ti,ab,kw
3. MeSH descriptor: [Electroencephalography] explode all trees
4. MeSH descriptor: [Magnetoencephalography] explode all trees
5. #1 AND (#2 OR #3 OR # 4)
6. MeSH descriptor: [Therapeutics] explode all trees
7. MeSH descriptor: [Psychiatric Somatic Therapies] explode all trees
8. MeSH descriptor: [Pharmaceutical Preparations] explode all trees
9. MeSH descriptor: [Heterocyclic Compounds] explode all trees
10. MeSH descriptor: [Organic Chemicals] explode all trees
11. MeSH descriptor: [Psychotherapy] explode all trees
12. MeSH descriptor: [Behavior and Behavior Mechanisms] explode all trees
13. ((therapeutic* or "brain stimulation" or tACS or tDCS or TMS or rTMS or ECT or LFMS or tPEMF or TBS or iTBS or cTBS or tPCS or tRNS or CES or "sensory stimulation" or modulat* or alter* or nicotin* or smoking or alcohol* or drug* or pharma* or music* OR neurofeedback or "neuro feedback" or "biofeedback" or "bio feedback" or meditat* or exercis* or physical activit* or relax* or task)):ti,ab,kw
14. (((brain* or cortex* or cortical* or crani* or transcrani* or magneti*) NEAR (stimulat* or electrostim* or "electro stimulation" or electrotherap* or "electro therap*"))):ti,ab,kw
15. (((auditory or visual* or nocicept* or sensory or somatosensor* or tactil* or light* or photic*)  NEAR (stim* or flicker))):ti,ab,kw
16. (#6 or #7 or #8 or #9 or #10 or #11 or #12 or #13 or #14 or #15)
17. #5 AND #16

Limit to trials (i.e. to exclude literature reviews)

**Unpublished Grey literature**

- The U.S. National Library of Medicine (ClinicalTrials.gov)
    - Search each term separately in ‘other terms’: alpha frequency, mu, frequency, dominant frequency, dominant peak, alpha peak
    - limit to ‘Adult (18-64)’
    - limit to ‘Accepts Healthy Volunteers’
    - limit to ‘With results’
- The System for Information on Grey Literature in Europe (opengrey.eu)
    - Search: (alpha frequency OR mu frequency OR dominant frequency OR dominant peak OR alpha peak) lang:"en"
    - Limit to ‘English’
- The New York Academy of Medicine Grey Literature Report (www.greylit.org)
    - Search each term separately: alpha frequency, mu, frequency, dominant frequency, dominant peak, alpha peak
- Preprint archive search OSF (https://osf.io/preprints/discover)
    - Search: (EEG OR MEG OR electroencephalogra* OR magnetoencephalogra*) AND ("peak alpha frequency" OR "dominant peak" OR "alpha peak" OR "mu frequency" OR "dominant frequency" OR "individual peak")
    - No limits
