## Supplementary Material 2 for "Can non-invasive brain stimulation modulate peak alpha frequency in the human brain? A systematic review and meta-analysis"

### PAF Systematic Review - EEG methodology

Start of Block: Default Question Block

Q1 Reviewer:

- ◯ SKM
- ◯ DS
- ◯ PS
- ◯ Other ________________________________________________

Q2 Article Title: (as seen on article)

________________________________________________________________

Q3 First three author's names: (as seen on the article, if more than 3, add et al)

________________________________________________________________

Q4 Article published date: (as seen on article)

________________________________________________________________

Q91 Please indicate what experiment you are reporting on: (if only one experiment within the article, please select "A")

- ◯ A
- ◯ B
- ◯ C
- ◯ D

End of Default Question Block

Start of Block: Block 1

*Each question within this block has Yes, Partially, No, Comment, and NA response options*

Q5 **Terminology and experimental breakdown:** Are all experimental events described consistently using COBIDAS lexicon?

Q6 **Terminology and experimental breakdown:** Was number of sessions, runs per session and trials/events per experimental condition collected consistently across participants and reported?

Q7 **Terminology and experimental breakdown:** Did article record that analyses were done in sensor or source space (or both)?

Q8 **Statistical Power:** Did the article detail any analysis performed a priori to justify the number of trials / participants?

Q9 **Participants:** Was recruitment, selection and sampling strategy recorded?

Q10 **Participants:** Did the study outline inclusion and exclusion criteria?

Q11 **Participants:** Demographics recorded?

Q12 **Participants:** Was information about written informed consent (or informed assent for pediatric participants) and the name of Institutional Review Board recorded?

Q13 **Stimulation/task parameters:** Was number of experimenters recorded?

Q14 **Stimulation/task parameters:** Were instructions (Task-related or not) provided to the participant recorded?

Q15 **Stimulation/task parameters:** Were stimulus properties recorded?

Q16 **Stimulation/task parameters:** Were calibration procedures recorded?

Q17 **Stimulation/task parameters:** Were structure and timing of the task (number of trials, ISI/SOA, temporal jitter, order of stimuli/conditions, counterbalance, etc) recorded?

Q18 **Behavioral Data:** If resting state data, did article indicate if the participant's eyes are open or closed? If open, did the article indicate whether a fixation point was used?

Q19 **Device:** Was MEG or EEG manufacturer, model, sensor specifications reported?

Q20 **Device:** Were details on additional devices used (manufacturer and make) for additional measures (behaviour or other) provided? If not applicable, please leave a comment as to why this is.

Q21 **Sensor type and spatial layout:** MEG: Were planar/axial gradiometers and/or magnetometers, and their number and locations reported? If not applicable, please leave a comment as to why this is.

Q22 **Sensor type and spatial layout:** Did the article report the EEG spatial layout: 10-20, 10-10 system, Geodesic, other? Did the article document number of electrodes? If layout is not conventional, did the article show a 2D map of electrode positions? If not applicable, please leave a comment as to why this is.

Q23 **Sensor type and spatial layout:** Were electrodes for EEG, EOG, ECG, EMG, skin conductance (electrode material, passive/active, other) recorded? If not applicable, please leave a comment as to why this is.

Q24 **Participant preparation and test room:** Were ambient characteristics and lighting (and if appropriate, empty room recording for MEG), detailed, and whether or not the recording room was shielded for EEG?

Q25 **Participant preparation and test room:** Was participant preparation recorded (skin preparation prior to electrode application, electrode application; participant degaussing, special clothing)?

Q26 **Impedance measurement:** Did the article report impedances for EEG/EOG/ECG/EMG electrodes, preferably digitally storing impedance values to the datafile, indicate timing of impedance measurement(s) relative to the experiment? If not applicable, please leave a comment as to why this is.

Q27 **Data acquisition parameters:** Were software systems used and system year for acquisition reported?

Q28 **Data acquisition parameters:** Were low- and high-pass filter characteristics reported?

Q29 **Data acquisition parameters:** Was sampling frequency reported?

Q30 **Data acquisition parameters:** Did the study report a continuous versus epoched acquisition?

Q31 **Data acquisition parameters:** For EEG/EOG/ECG/EMG/skin conductance: Did the article report reference and ground electrode positions? If not applicable, please leave a comment as to why this is.

Q32 **Sensor position digitization:** EEG/EOG: Was the method (magnetic, optical, other), manufacturer and model of the device used reported? If not applicable, please leave a comment as to why this is.

Q33 **Sensor position digitization:** MEG: Was the monitoring of head position relative to the sensor array, the use of head movement detection coils and their placement reported? If not applicable, please leave a comment as to why this is.

Q34 **Sensor position digitization:** In both MEG and EEG, was the time of digitization in relation to the experiment, and the 3D coordinate system reported?

Q35 **Workflow:** Did the article indicate in detail the exact order in which preprocessing steps took place?

Q36 **Software:** Did the article report which software and version were used for preprocessing and processing, and analysis platform?

Q37 **Software:** Was any in-house code used? If so, was it shared/made public? If no in-house code was used, please reflect this in a comment.

Q38 **Generic preprocessing:** Did the article report whether or not any downsampling of the data occurred?

Q39 **Generic preprocessing:** Did the article report if electrodes/sensors were removed, which identification method was used, which ones were deleted, and if missing channel interpolation is performed article indicated which method?

Q40 **Generic preprocessing:** Did the article specify detrending method (typically polynomial order) for baseline correction? If not applicable, please comment why.

Q41 **Generic preprocessing:** Did the article specify noise normalization method (typically used in multivariate analyses)? If not applicable, please comment why.

Q42 **Generic preprocessing:** If data segmentation is performed, article indicated the number of epochs per participant per condition? If not applicable, please comment why.

Q43 **Generic preprocessing:** Article indicated the spectral decomposition algorithm and parameters, and if these were applied before/after segmentation?

Q44 Did the article report artifacts?

*Skip To: Q53 If Did the article report artifacts? = N*

Q45 **Detection/rejection/correction of artifacts:** Did the article indicate what types of artifact are present in the data?

Q46 **Detection/rejection/correction of artifacts:** For automatic artifact detection, article described algorithms used and their respective parameters (e.g., amplitude thresholds)? If not applicable, please comment why.

Q47 **Detection/rejection/correction of artifacts:** For manual detection, the article indicated the criteria used with as much detail as needed for reproducibility? For not applicable, please comment why.

Q48 **Detection/rejection/correction of artifacts:** Did the article indicate if trials with artifacts were rejected or corrected? If using correction, did they indicate method(s) and parameters? If not applicable please comment why.

Q49 **Detection/rejection/correction of artifacts:** If trials/segments of data with artifacts have been removed, article indicated the average number of remaining trials per condition across participants (including minimum and maximum number of trials across participants)? If not applicable please comment why.

Q50 **Detection/rejection/correction of artifacts:** For resting state data, did the article specify the length of time of the artifact-free data? If not applicable, please comment why.

Q51 **Correction of artifacts using BSS/ICA:** Article indicated how many total components were generated, what type of artifact was identified and how, and how many components were removed (on average across participants)?

Q52 **Correction of artifacts using BSS/ICA:** Does the article display example topographies of the ICs that were removed?

Q53 **Filtering:** Was filter type (high-pass, low-pass, band-pass,band-stop, FIR, IIR) reported?

Q54 **Filtering:** Filter parameters: Was cut off frequency (including definition: e.g., -3 dB/half-energy, -6dB/half-amplitude, etc.), filter order (or length), roll-off or transition bandwidth, passband rippleand stopband attenuation, and filter delay (zero-phase,linear-phase, non-linear phase) reported?

Q55 **Filtering:** Only applicable in cases of two-pass filtering: Were causality and direction of computation reported (one-pass forward can only be reverse or forward, two-pass forward and reverse)? If not applicable, comment why.

Q56 **Re-referencing (for EEG):** Did the article report the digital reference and how this was computed?

Q57 **Re-referencing (for EEG):** Did the article justify choice of the re-reference scheme? If not applicable, comment why.

Q58 **ROIs:** Article reports how ROIs were determined, i.e. what was the mode of selection (e.g., a priori from literature or independent data)?

Q59 **ROIs:** Did the article report specific sensors/regions of interest, peaks, components, time and/or frequency window, source?

Q60 **Summary measures:** Did the article report how summary measures were obtained?

Q61 **Summary measures:** Did the article justify how the selection of dependent variables is unbiased (especially how the temporal and spatial ROIs were chosen)?

Q62 **Summary measures:** Did the article describe how peaks, components, and latencies were measured?

Q63 **Statistical analysis/modeling:** Article reported software and version used, and analysis platform reported?

Q64 **Statistical analysis/modeling:** Article reported model used including all regressors (and covariates of no interest)?

Q65 **Statistical analysis/modeling:** Article describes their check and report of statistical assumptions (e.g.,normality, sphericity)?

Q66 **Statistical analysis/modeling:** Article provides details on classification method and validation procedure?

Q67 **Statistical analysis/modeling:** Article notes method used for multiple comparisons correction and chosen level of statistical significance?

Q68 **Statistical analysis/modeling:** Article reports classifier used, the distance metric used and the parameters?

Q69 **Statistical analysis/modeling:** Article reports how chance level was determined?

Q70 **Statistical analysis/modeling:** Article details cross-validation scheme?

Q71 **Statistical analysis/modeling:** Article reports/justifies data reduction method and parameters if used (PCA, SVD, etc.)? Comment if not applicable.

Q72 **Nomenclature:** Article uses IFCN-sanctioned nomenclature?

Q73 **Time and frequency windows:** Did the article explicitly note the time and frequency windows?

Q74 **Statistical Results:** Did the article report the statistical values of analyses performed? For mass-univariate and multivariate approaches, did the article report minima and maxima of R-squared, z/t/F values?

Q75 **Statistical Results:** Did the article report raw effects (onset/offset in time, frequency, amplitude, power) and standardized effect sizes?

Q76 **Figures:** Did the article show waveforms or spectra of each condition and differences of interest (indicates clearly whether individual participant or group data are displayed)?

Q77 **Figures:** Did the article display waveforms with measures of error (confidence intervals / standard deviation over participants for the grand average, or over trials for individual participants)?

Q78 **Figures:** Did the article associate waveforms and spectra with topographic representations?

Q79 **Figures:** Did the article label all axes, reports units and if needed displays calibration bars (i.e., colorbar) with units?

Q80 **Figures:** Did mass-univariate and multivariate analysis show the full space of statistical results along with significant results?

**End of Block: Block 1**

---
